# Accurate Direct PCR with *Arabidopsis* and rice

**DOI:** 10.1101/2021.07.16.452406

**Authors:** Peter Lynagh, Paul Osuna-Kleist, Bohai Wang, Edgar Malagon, Maria Ximena Anleu Gil, Luca Comai

**Affiliations:** Department of Plant Biology, Davis, CA, USA; Genome Center, Davis, CA, USA; Innovative Genomics Institute, University of California, Berkeley, CA, USA

## Abstract

Plant sampling methods for PCR that avoid costly and time-consuming plant DNA purification can be accurate but have not been widely adopted, partly because the efficacy of the methods is unclear. Here, we introduce new sampling methods for Direct PCR in *Arabidopsis* and rice and compare them to previously published methods. CutTip, stabbing a pipette tip into a plant and depositing the tip of the pipette tip into the reaction buffer, yielded high accuracy for genotyping *Arabidopsis* and rice. This did not require visible tissue fragments in the reactions. We found that using short thin segments of fishing line, called Line-PCR, to be even more practical and highly accurate. Compared to traditional and other Direct PCR preparation methods, Line-PCR is faster, inexpensive, easier, uses less plastic and chemicals, and causes the least tissue damage. CutTip and Line-PCR require no modified PCR reagents. Line-PCR particularly lacks any complicated steps. Probably, the most difficult step in Line-PCR is a wipe of the tweezer between samples. These methods can help address the challenge of genotyping at different scales with high accuracy. CutTip and Line-PCR also simplify the application of PCR in the field.

## Introduction

The Polymerase Chain Reaction (PCR) continues to be one of the most powerful tools in molecular biology. However, PCR can frequently fail if starting with too few DNA molecules or impure DNA. Plant biologists tend to assume that plant DNA needs at least a rudimentary DNA purification step in order for PCR to have the desired success, *i*.*e*. high accuracy. Accuracy is defined by the combination of high sensitivity and high specificity, *i*.*e*. by low false negatives and low false positives. However, DNA purification can be expensive in money, time, and effort. The most popular DNA purification method entails high speed centrifugation steps in organic solvents such as chloroform and isopropanol followed by drying and dissolution of the DNA (Figure 1A, B). Embryophyta (land plants) is Earth’s most abundant phylum by mass and our primary food source (1). It is important to have efficient methods for PCR with plants.

**Figure 1.**
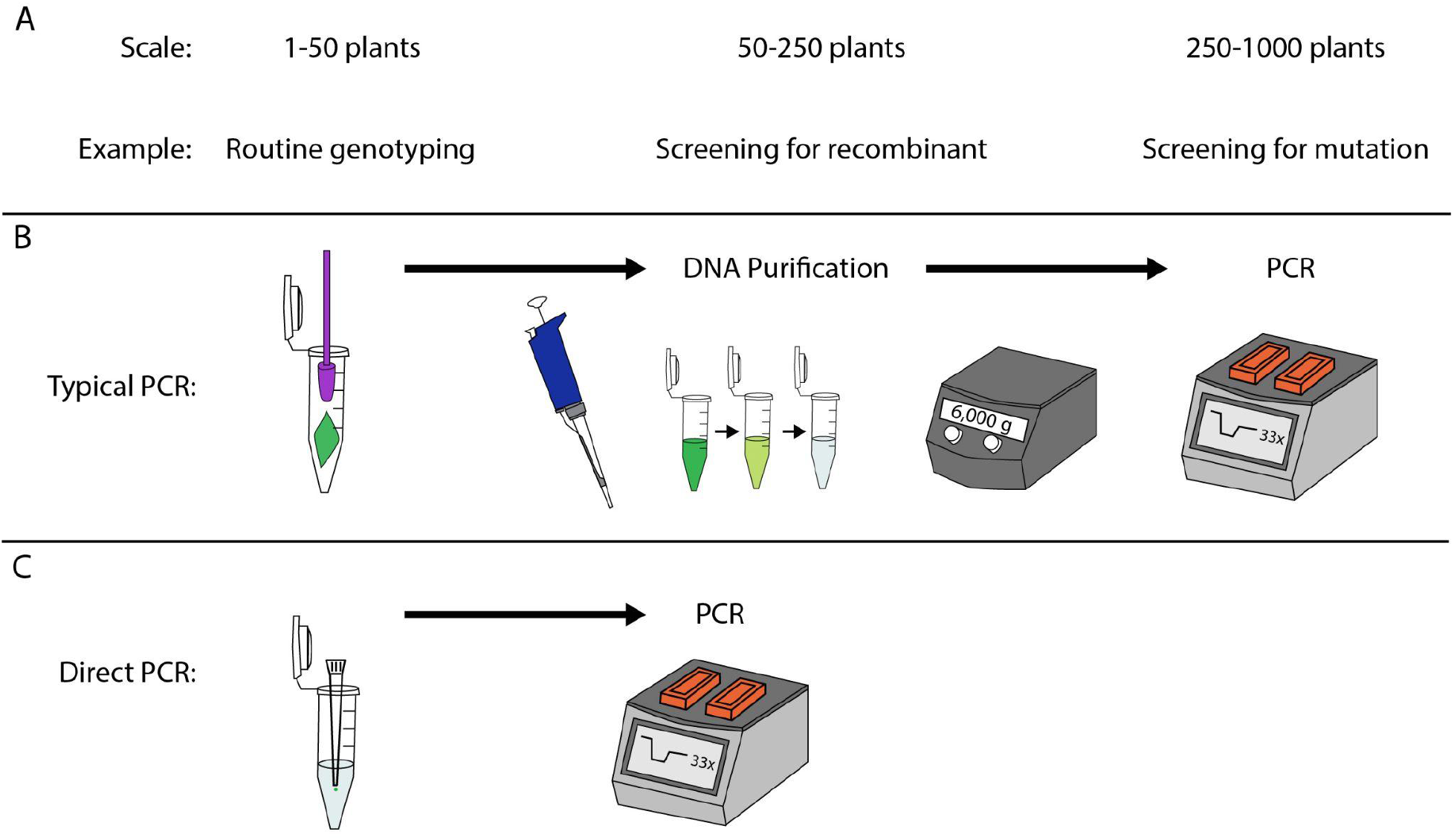
Plant Direct PCR helps overcome genotyping bottlenecks. A. Plant genotyping is common at different scales. B. Typical PCR uses a DNA purification step. C. Direct PCR skips DNA purification.

The DNA purification steps can be minimized or eliminated. Direct PCR is defined as PCR that does not include any DNA purification step in the protocol (Figure 1C). The strict definition of Direct PCR used in this article includes only methods in which impure DNA is directly transferred from the plant to the PCR reagents. Thus, Direct PCR can be relatively inexpensive, fast, and easy. Though not commonly used, Direct PCR has been shown to work in many plant species. Amplification by Direct PCR has been demonstrated in dozens of diverse plant species including liverworts, mosses, ferns, monocots and dicots (2); (3); (4); (5); (6); (7); (8).

Although plant Direct PCR clearly can work, one reason that plant Direct PCR has not become popular is that plant tissue can strongly inhibit PCR (3). In some species, tissue pieces as small as 1 mm.^2^ are inhibitory (7). Specialized plant Direct PCR buffers that lessen inhibition are commercially available, but we wondered if there might be a simpler way to achieve high plant Direct PCR accuracy.

One of the simplest Direct PCR methods is to use a scalpel or tweezers to pick a roughly 1 mm.^2^ piece of tissue and put it directly in the PCR reagents (9). It requires an attentive technician if tissue >1 mm.^2^ inhibits the reaction. One of the first and simplest plant Direct PCR methods described involves poking a leaf with a pipette tip, then pipetting the visible tissue directly into the PCR reagents (2), which we call the Berthomieu-Meyer (B.M.) method. The study noted that some of the reactions failed (2). The same method was attempted with smaller pipette tips that acquire smaller tissue, occasionally using a syringe needle to push the tissue out of the pipette tip improving the PCR sensitivity (3). Another study showed good sensitivity using wooden toothpicks to directly transfer DNA from the plant to the PCR reagents (5).

Although some studies on Direct PCR reported sensitivity, the true positive rate, we are unaware of any reporting specificity, the true negative rate. It is essential to report both sensitivity and specificity especially when cross-contamination is a possibility. Specificity can drop all the way to zero if care is not taken to avoid cross-contamination. This is not unique to Direct PCR. Most DNA purification methods are at risk of False Positive PCR reactions due to cross-contamination. We tested methods analyzing their sensitivity, specificity and accuracy using *Arabidopsis* and rice. To address shortcomings, we developed Direct PCR methods that we call CutTip and Line in which tissue is poked with a probe and dropped into water or the PCR reagents where the probe stays during the entire PCR. Our findings provide improvements that should benefit many studies.

## Materials and Methods

### Choosing loci and designing primers

The end goal is to measure the sensitivity and specificity of attaining bands that we assume contain a predetermined sequence of interest. As Direct PCR is not expected to have a higher sensitivity than traditional PCR, loci that do not amplify when using CTAB-purified genomic DNA were excluded. Eleven Col-0 and 8 L*er*-specific targets were amplifiable using CTAB-purified DNA and were used to measure Direct PCR with *Arabidopsis*. The 8 *Arabidopsis* L*er* primer pairs were chosen based on having smaller expected band sizes as compared to Col-0 (10). Though not completely randomized, we had never attempted amplification of 6/12 of the Col-0 loci and 7/8 of the L*er* loci. All except three *Arabidopsis* primers worked, mostly due to L*er* sequence not matching the reference. The 3 non-performing *Arabidopsis* primers were replaced with alternative primers that amplify the same locus, which we allowed because it is a common optimization step regardless of the method of PCR. For rice, sequences of interest were randomly selected from within segments of 1-5 megabases of Chromosomes 4-7. All chosen rice primer pairs worked well. In summary, we believe that our study used loci that are representative of what researchers typically encounter in *Arabidopsis* and rice.

Candidate primer sequences were chosen by manually looking in the upstream and downstream portions of the sequences of interest. Desired primer characteristics included mixing of all 4 nucleotides especially on the 3’ side, avoidance of repeats, a G or C at the 3’ end, 26-31 nucleotide length and 150-500 bp between primers of each pair. The “PCR Primer Stats” tool on www.bioinformatics.org was used to find and avoid primer self-annealing. Primers are listed in Supplemental Tables 1 and 2.

### Direct Tissue vs. CTAB

What we call “CTAB DNA” is Col-0 DNA that was purified using a CTAB protocol (11). Fresh tissue was frozen with liquid nitrogen and ground with a pestle. Five-hundred μL CTAB extraction buffer was added, then shaken. One-hundred fifty μL 24:1 Chloroform:Isoamyl Alcohol was added and shaken. After centrifuging at 4,000 xg for 5 minutes, the CTAB layer was transferred to a tube containing 200 μL isopropanol and shaken. After centrifuging at 8,000 xg for 5 minutes, the isopropanol was poured out and 400 μL 70 % ethanol was added. The 70 % ethanol was immediately poured out. The purified DNA was air dried at 37 °C for about 1 hour. The DNA was dissolved in nuclease-free water.

For the Direct Tissue PCR, tissue was simply picked with tweezers and then tapped into the tube. For estimated the size of the tissue, front and side images of each tube were taken using a dissecting microscope. Adobe Photoshop 2021 was used to roughly estimate the tissue Area using the Lasso tool and Image -> Analysis.

### Pipetting Direct PCR

Pipetting tissue Direct PCR was done as originally described (2, 3). For Arabidopsis, a 20 μL pipette tip was poked through a leaf, using a flat wooden toothpick or popsicle stick as a surface to press against. We avoided pressing onto the same spot on the popsicle stick more than once, but were able to do several pokes on each popsicle stick. The pipette tip was then placed onto the pipette and pipetted up and down 10 times in the PCR solution. For rice, a 1,000 μL pipette tip was poked through a leaf, using a popsicle stick as a surface. After removing the pipette tips, the reaction plate was sealed, briefly vortexed, briefly centrifuged and then the PCR reaction was started.

### CutTip PCR

Materials are shown in Supplemental Figure 2. A visual protocol can be viewed in Supplemental Movie 1. The CutTip method starts with the same steps as the Pipetting Direct PCR methods. The only difference is that, in CutTip, the tip of the pipette tip is cut off and dropped into the reaction well that already contains all of the PCR reagents. Common cosmetic scissors work satisfactorily, though we prefer push spring scissors due to ease of use. We use model HTS 144S7 4.5” Straight Stainless Steel Squeeze Scissors purchased from www.amazon.com. The CutTip stays in the reaction during the PCR and we never remove the CutTip. For *Arabidopsis* we use 20 μL pipette tips, and either a flat wooden toothpick or popsicle stick as a surface to press on. For accuracy measurement with rice we use T-First-CutTip with 1,000 μL pipette tips and only popsicle sticks as a surface, though a 20 μL tip works. For rice we used NT-CutTip with iteratively cut p20 pipette tips exactly like we did NT-CutTip with *Arabidopsis*. We alternated many real positive samples with negative controls that did not involve poking tissue but did receive a CutTip. For measuring specificity and accuracy, we alternated every positive plant with a negative plant so that we could get a minimally biased measure of cross-contamination. When measuring sensitivity but not accuracy nor specificity, we interspersed each set of 10 positive samples with 4 negative controls that received a CutTip but presumably no template DNA.

What we call “CutTip” involves iteratively cutting up the pipette tip so that 1-12 samples can be taken using a single pipette tip. In contrast, “First CutTip” involves using only the very first cut of a pipette tip, then discarding the pipette tip, meaning only 1 sample for each pipette tip.

Errors occur when learning how to do this technique. One error is not getting the CutTip into the liquid, which occurs more with the lightweight 20 μL tips than the heavier 1,000 μL tips. The scissors and tips can be angled and placed next to the liquid to get all CutTips into their liquid. Another source of error is the scissors causing cross-contamination between wells. To minimize cross-contamination, we use a double dip system to clean the scissors between each sample. The scissors are first dipped into a beaker of water, then wiped with a wet paper towel. The scissors are then dipped into a second tube of water, then wiped with a dry paper towel. Typically, the next sample can use the same beaker of water and paper towel. See Supplemental Table 3 for more troubleshooting guidance.

### Line-PCR

Thirty meters of 0.279 mm. diameter fishing line (such as Rio 0X clear PowerFlex fly fishing tippet) was cut with scissors into 1-4 mm. segments, which is enough for 10,000 reactions. The easiest way to cut the line is to hold several centimeters of line vertically and cut with small scissors, such as cosmetic scissors, moving the scissors upward after each cut. This averages roughly 4 seconds to make each cut, which was faster and easier than chopping. Pointed tweezers (such as cosmetic brand Tweezees) are used to pick up and grasp a line segment. The line is poked through a leaf once. The line is placed into the 2.5-5 μL PCR reagents or water. The tweezers are dipped into a beaker of clean water, then wiped with a soaking wet paper towel. The tweezers are dipped into a second beaker of clean water, then wiped with a dry paper towel. Repeat the process with a new line segment for the next reaction tube. Each paper towel and beaker of water can be used for several pokes. Each beaker of water can last for hundreds of pokes without causing cross-contamination. Regarding cross-contamination, the bigger concern is to consciously wipe the tweezer between each poke. Though done at room temperature, evaporation occurs. Keep most of the rows sealed to minimize evaporation. After all lines are in sealed tubes, centrifuge if necessary and start the PCR. The protocol can be viewed in Supplemental Movie 2.

Thin-Line-PCR is similar but the process is modified to make it easier to handle the thin line (such as Trouthunter 10X fly fishing tippet, 0.074 mm. diameter, which was used in this manuscript, or the firmer Hitena 9X multi-polymer XT 0.08 mm. diameter, which is our preferred thin-line at the time of manuscript submission), which is so thin that it is difficult to grasp, see and poke with. We do not pre-cut thin lines, because it is difficult to grasp and lift the thin segments from a dish. Instead, we grasp the thin line roughly 1 mm. from its end while it is still attached to the spool. Then, we use scissors to make 1 cut on the other side of the tweezer, preferably cut at an angle to give the thin line a sharper point for piercing the leaf. Making the thin line segment ∼4 mm. long can make it easier to see when confirming that it is in the reaction tube, but make sure that the poking end of the segment is submerged into the liquid. We use an optivisor to assist with seeing the thin line poke through the leaf. Besides these modifications for thin lines, the Thin-Line protocol is essentially the same as the Line-PCR protocol.

### PCR reagents and parameters

All PCRs were performed in 10 μL reactions on Bio-Rad 96-well thermal cyclers. Each reaction contained 5 μL water (Hyclone), 4 μL 2X GoTaq Green Master Mix and 1 μL 5 μM primer pair. The poked leaf DNA was added by either pipetting or a CutTip as described above. The only PCR parameters that varied were annealing temperature, elongation time and number of cycles (Supplemental Table 1). Annealing temperatures were chosen by NEB Tm Calculator and ranged 50-61 °C (version 1.13.0). Elongation time was 33-55 seconds, depending on amplicon length and preliminary PCRs using CTAB-purified DNA. Every PCR reaction was within these thermal cycler parameters:

Step 1: 120 seconds, 95 °C

Step 2: 15 seconds, 95 °C

Step 3: 30 seconds, X °C Step 4: Y seconds, 72°C

24-40 cycles of steps 2-4

Step 5: 2:00, 72 °C

Step 6: Finished, room temperature

We have noticed occasional batch variation of primer orders. We recommend increasing the number of cycles if the amplicon bands are faint. Fifty cycles are possible while maintaining a good specificity.

### Sensitivity, Specificity and Accuracy

For *Arabidopsis*, the unit of measurement was the presence or absence of a band at an expected band size. Depending on whether the band was expected in a reaction, the result of each reaction could be classified as a True Positive (TP), True Negative (TN), False Positive (FP) or False Negative (FN). The following formulas were used for sensitivity, specificity and accuracy:

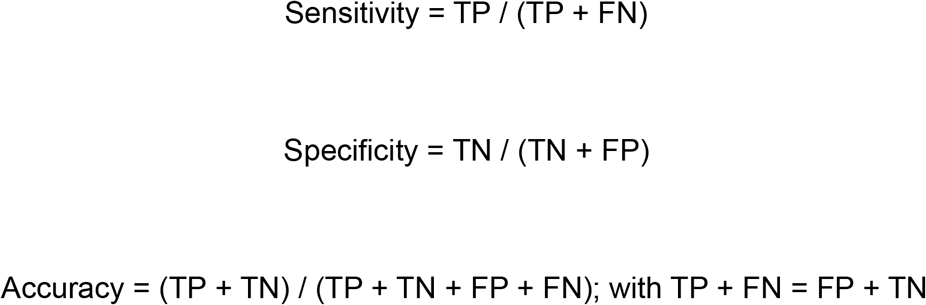

Directly measuring specificity requires having a clear and relevant definition of TN. We believe this was achieved for *Arabidopsis* by defining TN as a PCR band that is expected from L*er* but not expected from the closely related ecotype Col-0. Alternating each real Negative poke (Col-0) with a real Positive poke (L*er*) was essential for correct measurements of specificity.

For our rice population, there were technical and feasibility hurdles regarding the measurement of specificity. We had only one cultivar of rice on hand, and therefore we could not measure specificity and accuracy in rice in the same way that we did with Arabidopsis Col-0 and L*er*. Due to the invariable presence of the TP band when using the rice variety Kitaake and primer pairs designed using the Kitaake reference genome sequence, we lack the ability to do a simple specificity test. The rice loci were random and provided few suitable restriction sites. Even if we successfully digested or sequenced every band, the samples still would not fit the strict definition that TN does not produce the band to begin with. Due to such problems, we decided to define and use a Specificity Index for rice rather than directly measuring specificity. We define the Specificity Index as the presence *vs*. absence of PCR bands corresponding to DNA sizes different from the TP band. Thus, the Specificity Index is a measure of noise bands. Vague electrophoretic DNA smears and primer dimer products were not counted in the Specificity Index.

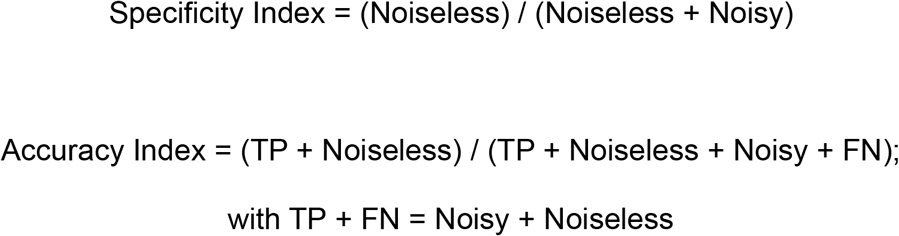

### Plant growth

*Arabidopsis* was grown on Sunshine Mix #1 soil fertilized by UC Davis Controlled Environment Facility (CEF) nutrient water. The light cycle was 14 hours light and 10 hours dark. The temperature was 17-23 °C. For measurement of sensitivity, specificity and accuracy, genotyping was performed using the healthiest leaves at the 6 leaf to first bolt stage. Rice was grown on a 16 hour light cycle with Sunshine Mix #1 soil fertilized by UC Davis CEF nutrient water. The temperature was a constant 24 °C. Rice genotyping was performed using the healthiest leaves on 1-2 month old plants.

## Results

To get a working baseline, we attempted Direct PCR by picking tissue with tweezers and putting the tissue directly into the PCR reagents (Figure 2A-D). While this has been done before, we replicated it because it is simple and intuitive. However, as reported previously, this method sometimes fails (Figure 2C). Even adding CTAB-purified template DNA to the tubes with tissue did not increase the sensitivity (Figure 2D). This fits with the previous findings that *Arabidopsis* PCR reactions containing tissue fail due to PCR inhibition caused by excess plant tissue. In addition to the low sensitivity, it took effort to repeatedly pick only small tissues (∼1 mm.^2^), transfer it into the solution and clean the tweezers. Therefore, we deemed this approach unpractical.

**Figure 2:**
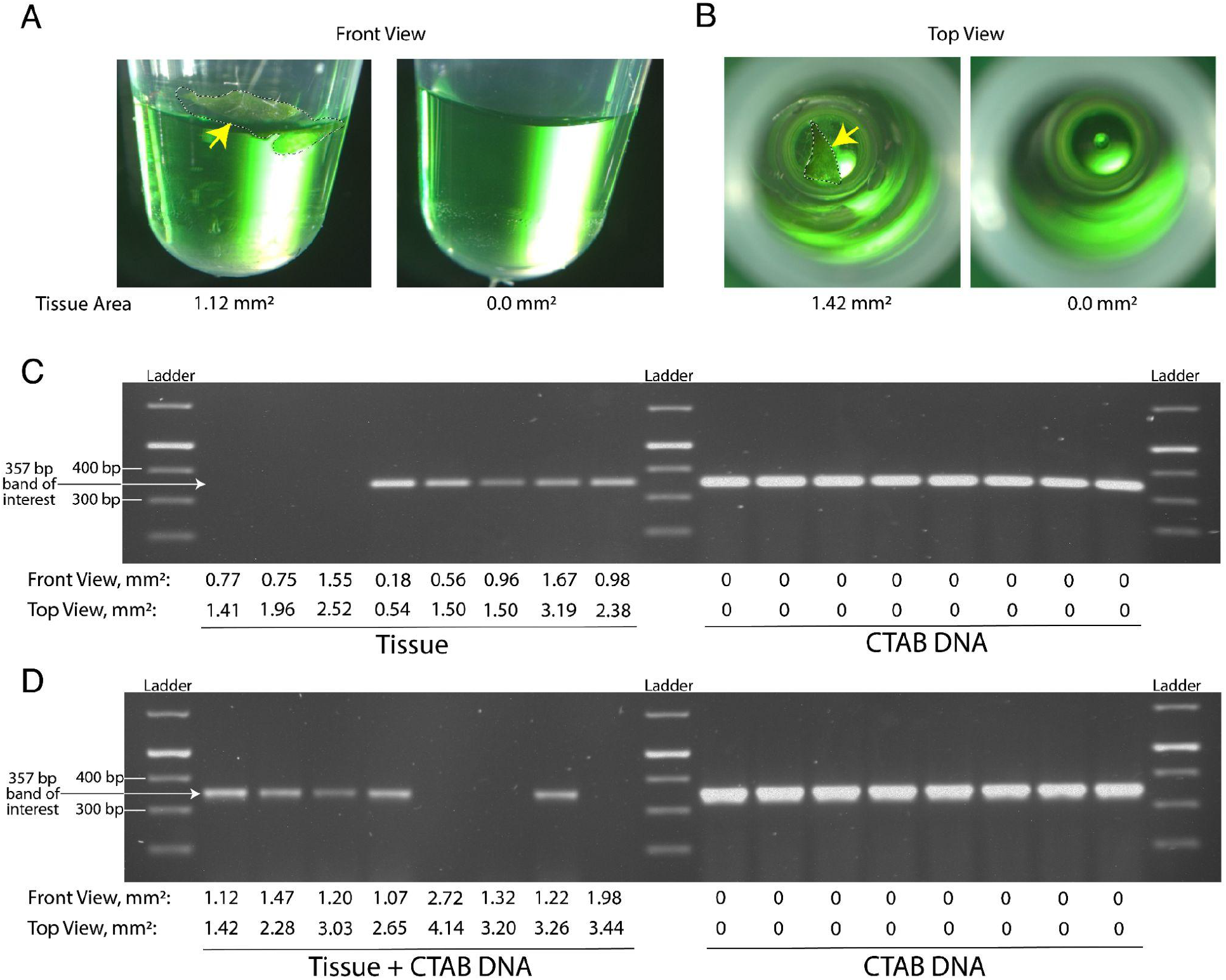
*Arabidopsis* tissue inhibits PCR. A. Example Front View of reactions just before PCR. Area within the tissue border is measured. Yellow arrow points to tissue. B. Example Top View of reactions just before PCR. Area within the tissue border is measured. C. Result of 16 PCR reactions, with an expected band size of 357 bp. The left 8 reactions contained Col-0 tissue but not CTAB DNA. The right 8 reactions contained CTAB DNA but no tissue. The numbers are estimates of the tissue size. D. All 16 PCR reactions contained the exact same reagents including a Col-0 primer pair and CTAB-purified Col-0 genomic DNA. The only difference is the left 8 reactions also received a piece of Col-0 tissue.

In preliminary work examining leaf poking methods for Direct PCR, we found that while wooden toothpicks often worked, the wood quickly absorbed the PCR liquid (Supplemental Figure 1A, B). We therefore rejected the use of wooden toothpicks for poking leaves. We found that plastic toothpicks often worked. We poked a leaf with a plastic toothpick, tap or vortex the plate with the plastic toothpicks and finally remove the plastic toothpicks before starting the PCR (Supplemental Figure 1C). When using plastic toothpicks like this, PCR sensitivities were 87.5% and 56.3% for 2 control primer pairs. We noticed that cutting the tip of the plastic toothpick and leaving it in the PCR reaction during the PCR potentially increased the sensitivity. The one drawback was that cutting plastic toothpicks took much physical effort due to the hardness of each toothpick (Supplemental Figure 1D). Thus, cutting toothpicks was rejected as a method. A softer material was needed. Due to the relative softness of pipette tips, we found it much easier to cut pipette tips. Overall, our preliminary data suggested that Direct PCR could immediately improve our screens for mutations in plants, but the PCR accuracy of the most promising methods had to be measured.

We compared four Direct PCR methods that involved poking an *Arabidopsis* leaf with a pipette tip (Table 1; Figure 3A; Supplemental Table 1). We reasoned that some False Negatives could result from insufficient release of plant material into the reaction. One solution is transfer of a visible piece of tissue from the pipette tip to the reaction mix, such as the B.M. method (2). Another was inspired by our preliminary experience with plastic toothpicks (Supplemental Figure 1C-1.1D; Figure 3B-I). We noted that after a poke in which visible tissue is not harvested, cell material is still present as a microscopic smear adhering on the pipette tip walls (Figure 3G, I). This material might not be released by simple dipping and pipetting. We therefore cut and released into the reaction mix a 1-3 mm. section of the pipette tip, which is the defining step of CutTip.

**Table 1:** The seven Direct PCR methods that are compared in this analysis. Method prefix “T” means tissue on tip is visible by eye. Method prefix “NT” means no tissue on tip is visible by eye. Y = yes. N = no. N/A = not applicable.

| Method | Pipetting, CutTip or Line | Tissue visible by eye? | Tissue size | Iterative Cutting? | Samples per pipette tip |
| --- | --- | --- | --- | --- | --- |
| Berthomieu-Meyer (B.M.)(2, 3) | Pipetting | Y | Small | N/A | 1 |
| NT-Pip | Pipetting | N | Absent or microscopic | N/A | 1 |
| T-CutTip | CutTip | Y | Small to large | Y | 2-12 |
| NT-CutTip | CutTip | N | Absent or microscopic | Y | 2-12 |
| T-First-CutTip | CutTip | Y | Small | N | 1 |
| Line | Line | N | Absent or microscopic | N/A | N/A |
| Thin-Line | Line | N | Absent or microscopic | N/A | N/A |

**Figure 3:**
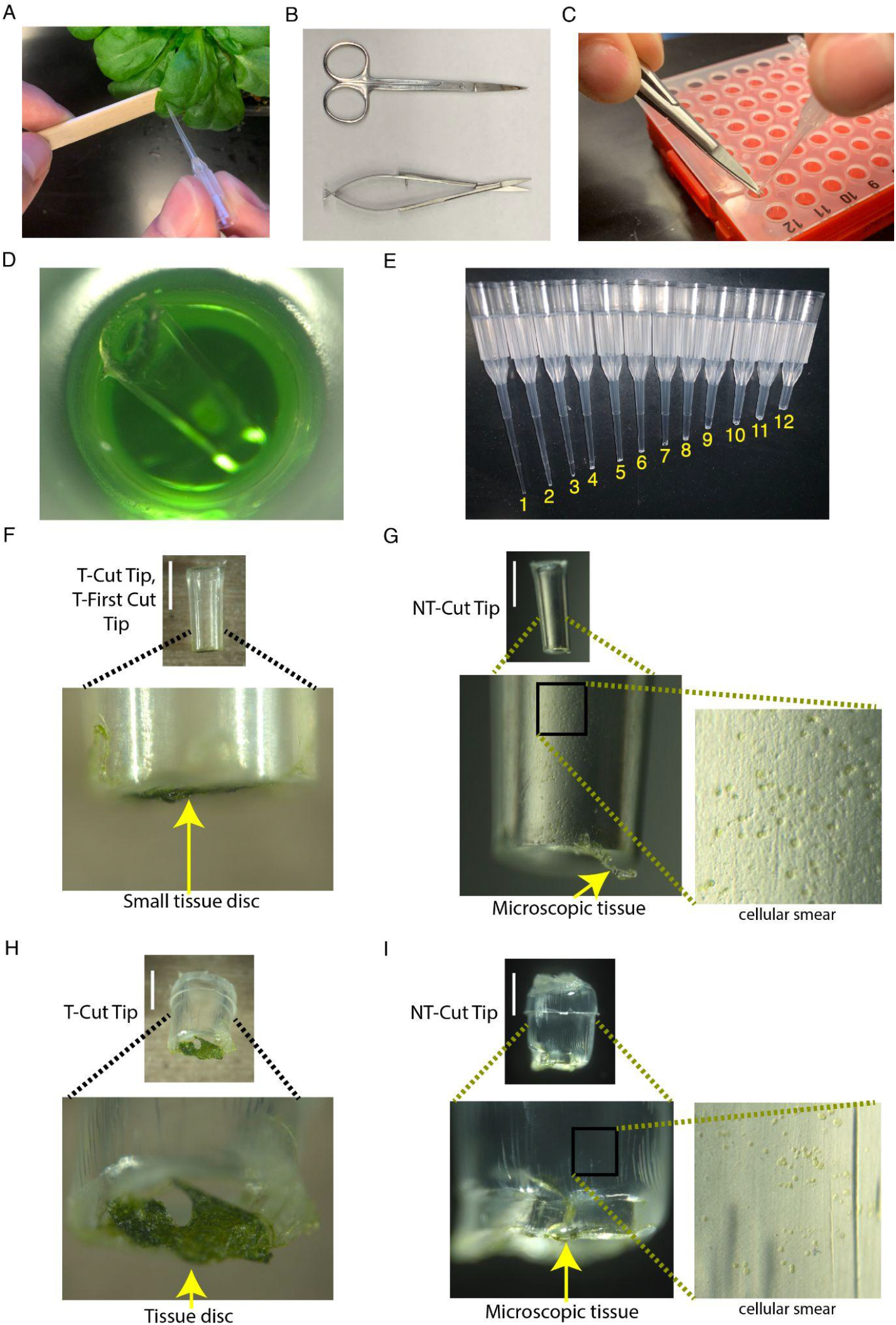
CutTip PCR. A. Poking an *Arabidopsis* plant with a pipette tip. B. Scissors that work for CutTip. The top scissors are easy to find. The bottom scissors work faster. C. Cutting and dropping the tip into the reaction reagents. D. Cut 20 μL tip in a 10 μL reaction. E. A single pipette tip can be iteratively cut with different samples taken with each cut. F. Typical T-CutTip or T-First-CutTip. G. Typical NT-CutTip. H. Typical T-CutTip. I. Typical NT-CutTip. Panels F-G correspond to location of cut #1 in Panel E. Panels H-I correspond to location of cut #10 in Panel E. F-I White Bar Scale = 1 mm.

As a result, two of the four tip-based Direct PCR methods involved cutting the tip of the pipette tip and leaving it in the reaction (Figure 3B, C and D; Supplemental Movie 1). We created this CutTip method in an effort to ensure sufficient DNA template is in the reaction and to increase sensitivity. The basic CutTip method involves iteratively cutting up the tip, with each cut releasing a different sample into a different well. If one is careful to avoid cross-contamination, it is possible to get 12 samples from 12 cuts of a single pipette tip (Figure 3E). The other factor used to differentiate the methods was either rejecting or selecting the presence of visible pieces of tissue on the tip after poking (Figure 3F, G, H and I). We prefix names of methods that have no visible tissue “NT” (Table 1). If an operator aims to obtain visible tissue (methods prefixed “T” as in T-CutTip), the tissue size increases as cuts are made further up the pipette tip (Figure 3E, F and H) eventually inhibiting the reactions and thus lowering sensitivity.

To measure sensitivity, we tested for the appearance of the PCR product in 20 technical replicates for each of 16 primer pairs resulting in 320 total reactions for each method at each specified number of cycles. At 24 PCR cycles, NT-Pip (meaning pipetting, no visible tissue) had the lowest sensitivity, likely due to suboptimal quantity of template DNA (Table 1; Figure 4A). T-CutTip had the highest sensitivity. Then, we repeated the experiment using 33 cycles. At 33 cycles, all four methods reached a fairly high sensitivity, though NT-CutTip and B.M. performed best with sensitivities of 97.9% and 97.3% respectively. We attributed the slightly imperfect sensitivity to the tip not sinking in the reaction liquid when using NT-CutTip. When using the B.M. method, imperfect sensitivity was attributed to failure of tissue transfer to the solution. We hypothesized that using only the first cut of a pipette tip with visible tissue (and discarding the rest of the pipette tip), called T-First-CutTip, would address both above problems and the tissue inhibition problem. T-First-CutTip includes only the very first cut of a tip, whereby the visible but small tissue can contact the solution. Thus, T-First-CutTip has two chances for template DNA to come into contact with the solution: 1) the cellular smear and 2) the visible small tissue. Using 8 different primer pairs, T-First-CutTip had a slightly higher average sensitivity than NT-CutTip, with 100% and 99.0% sensitivity respectively (Figure 4B). Either method has sufficient sensitivity for most experiments.

**Figure 4.**
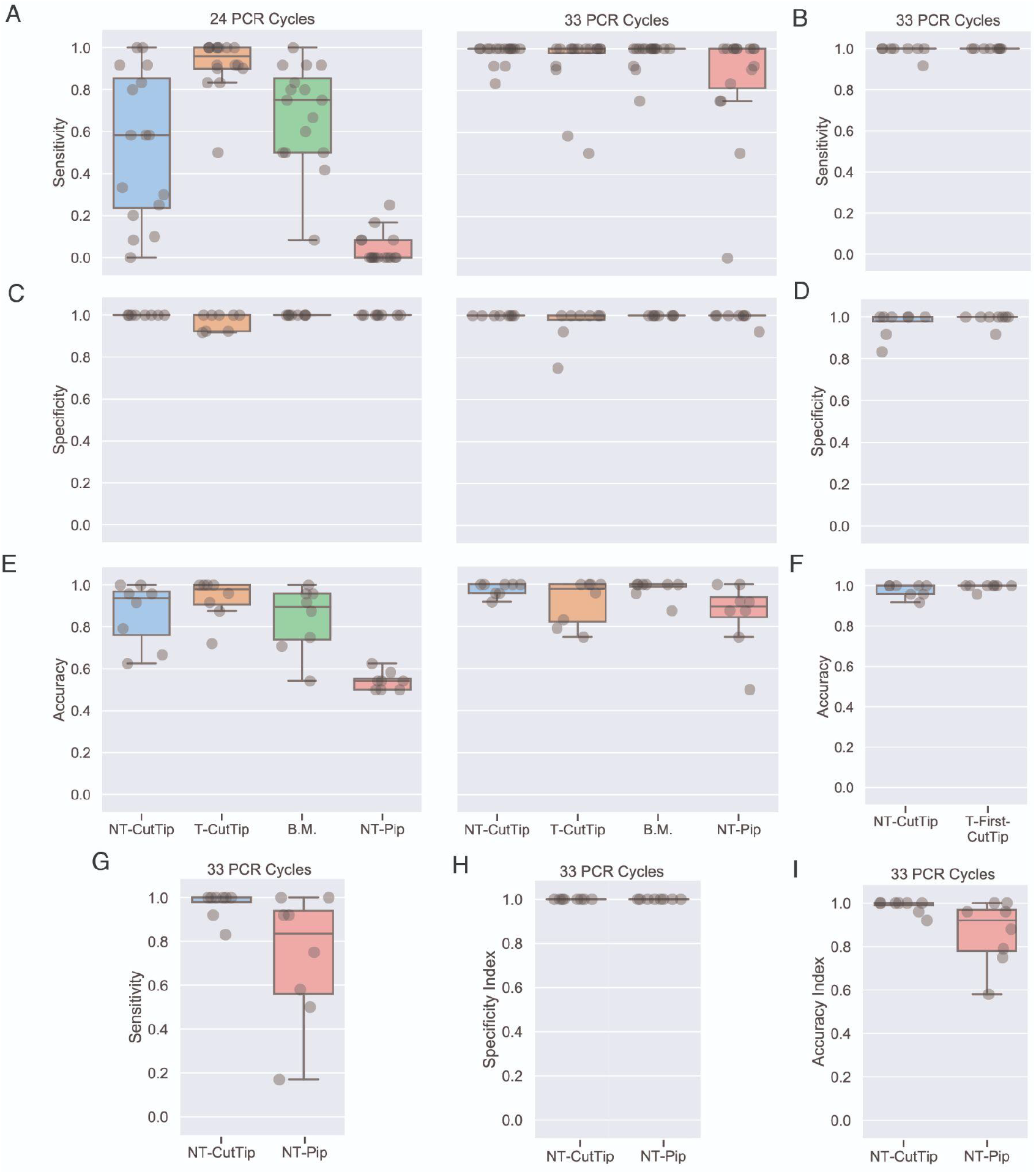
Accuracy of Direct PCR using *Arabidopsis* and rice. A.-F. Accuracy using *Arabidopsis*. A. Sensitivity using 4 Direct PCR methods. Each dot represents 1 of 16 primer pairs. B. Sensitivity after lessons from panel A. C. Specificity using 4 Direct PCR methods. Each dot represents 1 of 8 primer pairs. D. Specificity after lessons from panel C. E. Accuracy using 4 Direct PCR methods. F. Accuracy after lessons from panel E. G.-I. Accuracy using Rice. G. PCR Sensitivity Index of NT-CutTip and NT-Pip. H. PCR Specificity Index of NT-CutTip and NT-Pip. I. PCR Accuracy Index of NT-CutTip and NT-Pip.

In conclusion, B.M., T-First-CutTip and NT-CutTip provide high sensitivity. We next addressed specificity. To see how well Direct PCR can avoid amplifying incorrect sequences and cross-contamination, for each method we tested PCR amplification in 12 technical replicates for each of 8 primer pairs that are not expected to produce a product. We alternated pokes of real positives (Landsberg *erecta*, also called L*er*) and real negatives (Col-0). Overall, the specificity was high (Figure 4C and D).

The overall lack of bands of incorrect size in these PCRs suggest that the imperfect specificities were due mostly to template cross-contamination from the cut with a L*er* poke to the cut with a Col-0 poke, *i*.*e*. by carry over of template to the next cut segment. We hypothesized that T-First-CutTip would solve this problem because that method has no iterative cuts, meaning that each pipette tip is used for only one poke and the rest of the pipette tip is discarded. Indeed, T-First-CutTip had higher specificity than NT-CutTip, with 99.0% and 96.9% respectively (Figure 4D). Thus, NT-CutTip had 100% specificity in its first trial and 96.9% specificity in its second trial (Figure 4C and D). We believe the lower specificity of NT-CutTip in the second trial was due to the leaves being unusually plump and curvy that week. Thus, the properties of the tissue being sampled should be evaluated with common sense when using iterative pipette tip cutting.

Accuracy, measured as (TP+TN)/(TP+TN+FP+FN), is a summary composite of both sensitivity and specificity. A high accuracy means that plants that do not contain the DNA of interest do not yield the DNA band of interest, and plants that do contain the DNA of interest do yield the DNA band of interest. In other words, a high accuracy means the results contain a high proportion of True and low proportion of False. Based on their sensitivities and specificities, it is not surprising that NT-CutTip, B.M. and T-First-CutTip had the highest accuracies. B.M. and NT-CutTip both had 97.9% accuracy (Figure 4E and F). When compared to T-First-CutTip, T-CutTip again had accuracy 97.9% while T-First-CutTip had accuracy 99.5%. Practicality and accuracy are the critical criteria for method choice. While preparing the above 2,597 PCR reactions, we found that NT-CutTip had the fewest practical drawbacks of the five pipette tip methods (Table 2). Overall this suggests NT-CutTip and T-First-CutTip are the better methods for high-throughput genotyping, though B.M. has the same accuracy as NT-CutTip.

**Table 2:**
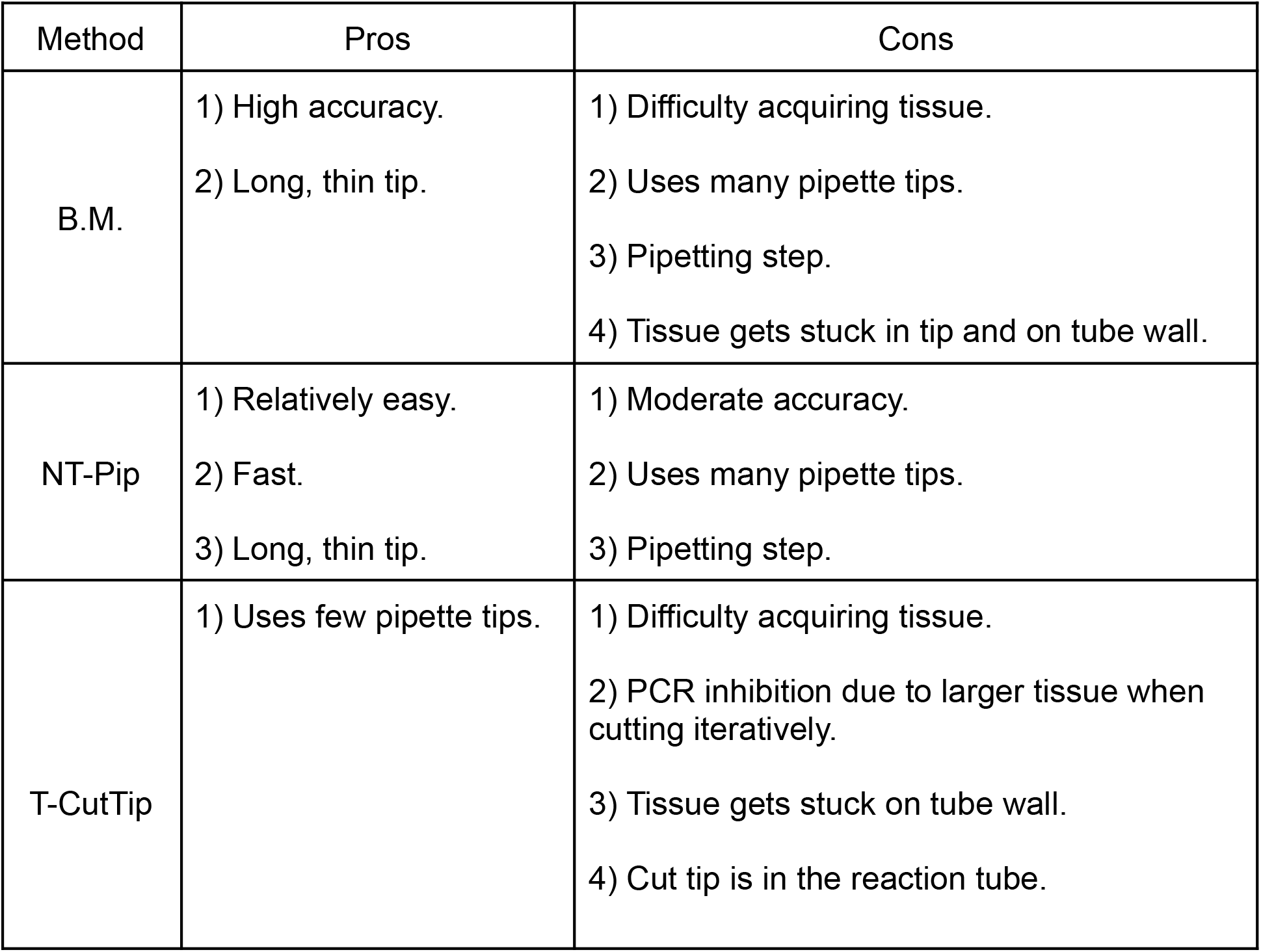
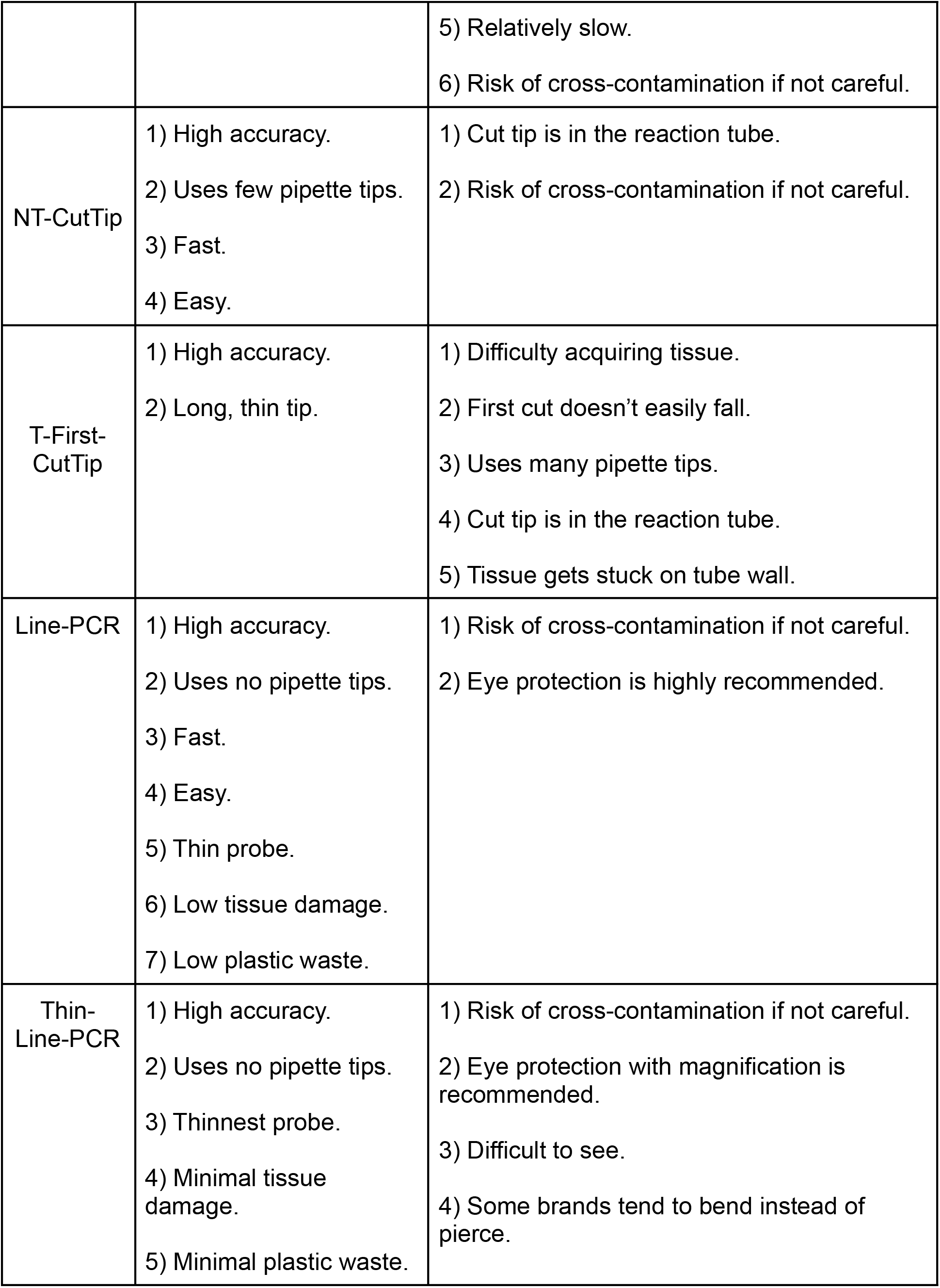
Pros and cons of sampling methods.

| Method | Pros | Cons |
| --- | --- | --- |
| B.M. | 1) High accuracy.<br>2) Long, thin tip. | 1) Difficulty acquiring tissue.<br>2) Uses many pipette tips.<br>3) Pipetting step.<br>4) Tissue gets stuck in tip and on tube wall. |
| NT-Pip | 1) Relatively easy.<br>2) Fast.<br>3) Long, thin tip. | 1) Moderate accuracy.<br>2) Uses many pipette tips.<br>3) Pipetting step. |
| T-CutTip | 1) Uses few pipette tips. | 1) Difficulty acquiring tissue.<br>2) PCR inhibition due to larger tissue when cutting iteratively.<br>3) Tissue gets stuck on tube wall.<br>4) Cut tip is in the reaction tube. |
|  |  | 5) Relatively slow.<br>6) Risk of cross-contamination if not careful. |
| NT-CutTip | 1) High accuracy.<br>2) Uses few pipette tips.<br>3) Fast.<br>4) Easy. | 1) Cut tip is in the reaction tube.<br>2) Risk of cross-contamination if not careful. |
| T-First-CutTip | 1) High accuracy.<br>2) Long, thin tip. | 1) Difficulty acquiring tissue.<br>2) First cut doesn't easily fall.<br>3) Uses many pipette tips.<br>4) Cut tip is in the reaction tube.<br>5) Tissue gets stuck on tube wall. |
| Line-PCR | 1) High accuracy.<br>2) Uses no pipette tips.<br>3) Fast.<br>4) Easy.<br>5) Thin probe.<br>6) Low tissue damage.<br>7) Low plastic waste. | 1) Risk of cross-contamination if not careful.<br>2) Eye protection is highly recommended. |
| Thin-Line-PCR | 1) High accuracy.<br>2) Uses no pipette tips.<br>3) Thinnest probe.<br>4) Minimal tissue damage.<br>5) Minimal plastic waste. | 1) Risk of cross-contamination if not careful.<br>2) Eye protection with magnification is recommended.<br>3) Difficult to see.<br>4) Some brands tend to bend instead of pierce. |

Next, we measured Direct PCR accuracy with rice. Sensitivity of Direct PCR was measured in rice using NT-First-CutTip and p1000 tips, though p20 tips also work (Figure 4G). In an effort to ensure our results would translate to other researchers’ rice genotyping, 8 random target loci were chosen in the first few megabases of chromosomes 4-7. The average sensitivity of NT-Pip in rice was 73.0% (Figure 4H). The average sensitivity using NT-First-CutTip was 96.9%. Because we had only one cultivar of rice on hand, we could not measure the specificity and accuracy in the way that we did with Arabidopsis Col-0 and L*er*. Instead of directly measuring specificity and accuracy, we measured a Specificity Index and Accuracy Index as explained in Materials and Methods. Any bands that are not at the expected position in relation to the DNA ladder cause the Specificity Index and Accuracy Index to decrease in the same way that a False Positive decreases specificity and accuracy. According to our measurements, both rice techniques had an average Specificity Index of 100%. The average Accuracy Index was 86.5% for NT-Pip and 98.5% for NT-First-CutTip (Figure 4I; Supplemental Table 2). Thus, NT-CutTip is an accurate Direct PCR method for rice genotyping.

Though pipette tips as probes work well, we felt that the procedure could be further simplified. We developed and tested the following methodology using Arabidopsis. We tried fishing line cut to segments of 1-3 mm. length, picking it up with tweezers, poking it into the leaf once, then dropping the line segment into the PCR reagents (Figure 5A-B, Supplemental Movie 2). We call this Line-PCR. Preliminary data suggested that various fishing line brands, materials and diameters work well. For measuring accuracy of Line-PCR, we chose 0.279 mm. diameter clear line (brand Rio 0X clear PowerFlex fly fishing tippet) because it causes little tissue damage while being thick enough to see and handle conveniently. Using five primer pairs, we found that Line-PCR had an average of 100% sensitivity, specificity, and accuracy (Figure 5C-E). As with CutTip, the high specificity hinged on cleaning the tweezers thoroughly between pokes. Though present, some of the bands were faint at 33 cycles, so we suggest doing 40 cycles if the amplification efficiency is low.

**Figure 5:**
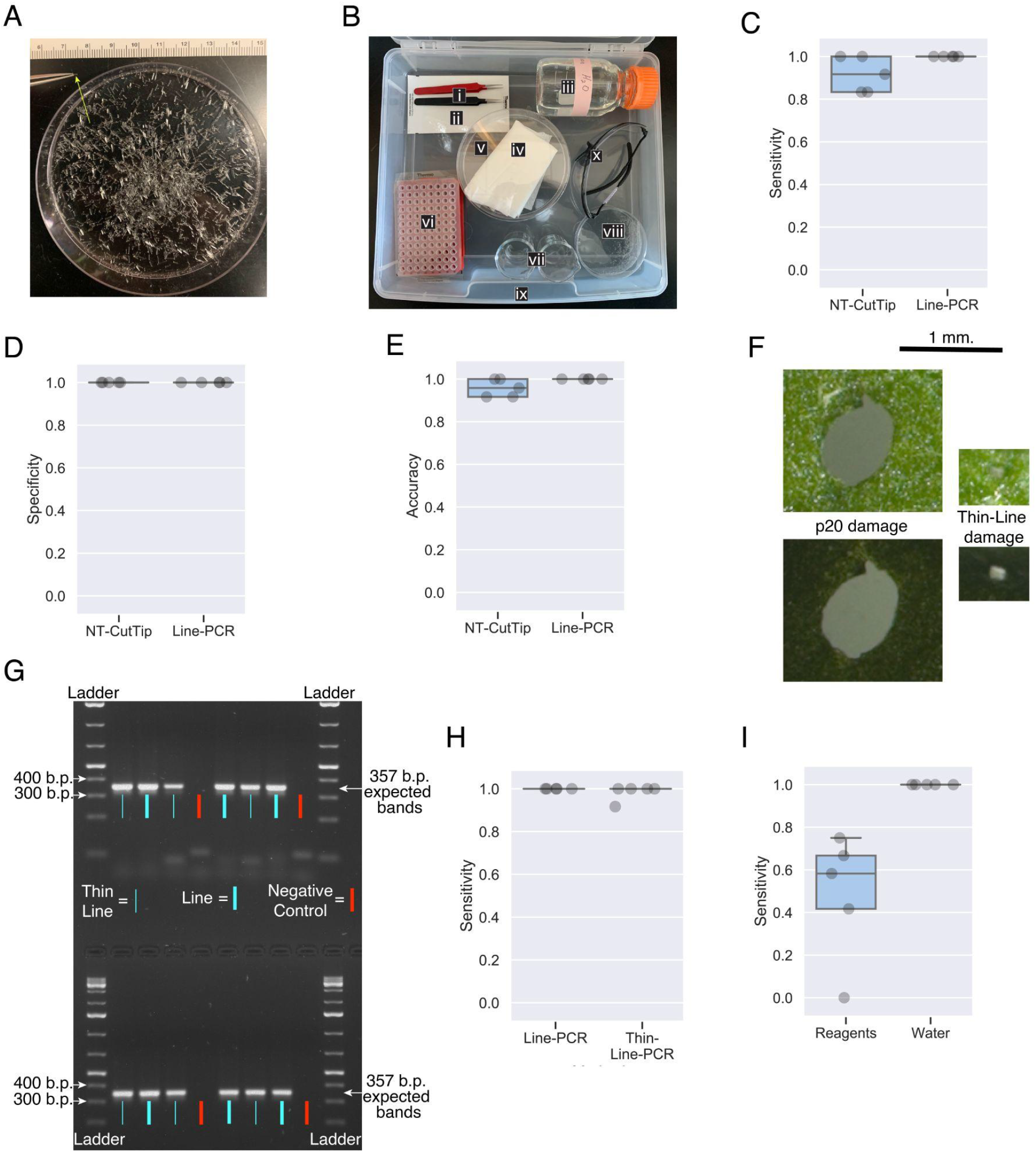
Line-PCR in *Arabidopsis*. A. Line segments sufficient for >2,000 reactions. Tweezers are holding one piece. Scale is millimeters. B. Example kit for Line-PCR. i. Two tweezers. ii. PCR sealing tape or caps. iii. Clean Water. iv. Paper towels. v. Clean dish. vi. Sealed PCR tubes containing all water or liquid reagents (not stored with kit). vii. Two beakers. viii. Line segments (the same dish shown in panel A). ix. Container for kit. x. Eye protection. C. Sensitivity of NT-CutTip versus Line-PCR. Each dot represents 1 of 5 primer pairs. D. Specificity of NT-CutTip versus Line-PCR. E. Accuracy of NT-CutTip versus Line-PCR. F. Hole in a leaf caused by a poke with a pipette tip versus a thin line (0.074 mm. diameter). The bottom and top photos are of the same holes. The bottom photo has higher contrast so that the hole from the thin line is visible. G. Example of PCR products when using line versus thin line to poke a plant to acquire template. H. Sensitivity of Line-PCR versus Thin-Line-PCR. Each dot represents 1 of 5 primer pairs. I. Stability of line poke samples. The sampling line was dropped into water (right) or into the complete PCR mix (left) and stored at room temperature for 24 hours. Then, master mix and primers were added to the water-only tubes and the PCRs were started.

We wondered if thinner line would be able to pierce a leaf and, if it could pierce, would the thin line have a high PCR sensitivity? The thinnest line that we could find is 0.074 mm. diameter. We found that the 0.074 mm. thin line does pierce leaves, though not as quickly as the normal (0.279 mm. diameter) line. The hole in the leaf caused by the poke is typically proportional to the diameter of the probe, so the thin line causes the smallest damage of all tested probes (Figure 5-F). We found that Thin-Line-PCR has a high sensitivity, with only one false negative in 60 reactions (Figure 5G-H). It is unclear if the false negative was caused by difficulty seeing that the thin line poked through the leaf or caused by the thin line not picking up enough template DNA during the poke. An example kit for Thin-Line-PCR includes an optivisor for improved visibility of the thin line while poking, and we found it easier to pick-up the thin line while it is still attached to the spool because it was difficult to pick-up the thin line from a dish (Supplemental Figure 4). We suggest Line-PCR for quick everyday genotyping. We suggest Thin-Line-PCR when there is a reason, such as minimizing tissue damage or poking a seedling. With a little practice, Thin-Line-PCR also works well for quick everyday genotyping.

Another concern is practicality and accessibility. We used a common PCR Master Mix with no modifications. All PCR reactions were prepared at room temperature. Regarding stability at room temperature, we prepared the PCR reactions and walked to and from the growth chambers (3 km) at room temperature for a total of about 2 hours before the PCR plates were put in the thermal cycler for Figures 4 and 5C-E. This suggests that Direct PCR can be prepared in the greenhouse or in the field. Most of the Direct PCR materials can be kept in a kit that can be taken to or stored with the plants, enabling PCR preparation and possibly PCR completion in the field (Figure 5B, Supplemental Figures 2 and 4).

We noticed that the PCR efficiency and sensitivity substantially drops after 2 hours of the probe being in the reagents at room temperature. We wondered if putting the cut tip or line into water instead of into the full PCR mix would allow for more time before putting the reactions into the thermal cycler. This is important because there is much variation in poking time depending on various parameters, such as the labeling of plants to poke and the speed of different technicians. To test if it is more stable to add line to water or to the PCR mix, we added line to both and waited 24 hours at room temperature before adding freshly thawed PCR mix to the line in water. We found that adding the line to water was far more stable for 24 hours as indicated by its 100% sensitivity *versus* 48% sensitivity when the line was added directly to the PCR reagents 24 hours before starting the PCR (Figure 5I). The only drawbacks to adding the line to water is the use of one small pipette tip for each reaction when pipetting the PCR mix and being careful to not accidentally remove the line while pipetting. The benefits of adding the line to water outweigh the drawbacks, so we suggest adding the line or cut tip to water if the poking process is expected to take longer than an hour. Integrating all that we learned, we recommend the Line-PCR protocol that is summarized in Figure 6. In summary, Line-PCR worked well on arabidopsis, as well as two other species whose results are not shown, potato (*Solanum tuberosum*) and camelina (*Camelina sativa*).

**Figure 6:**
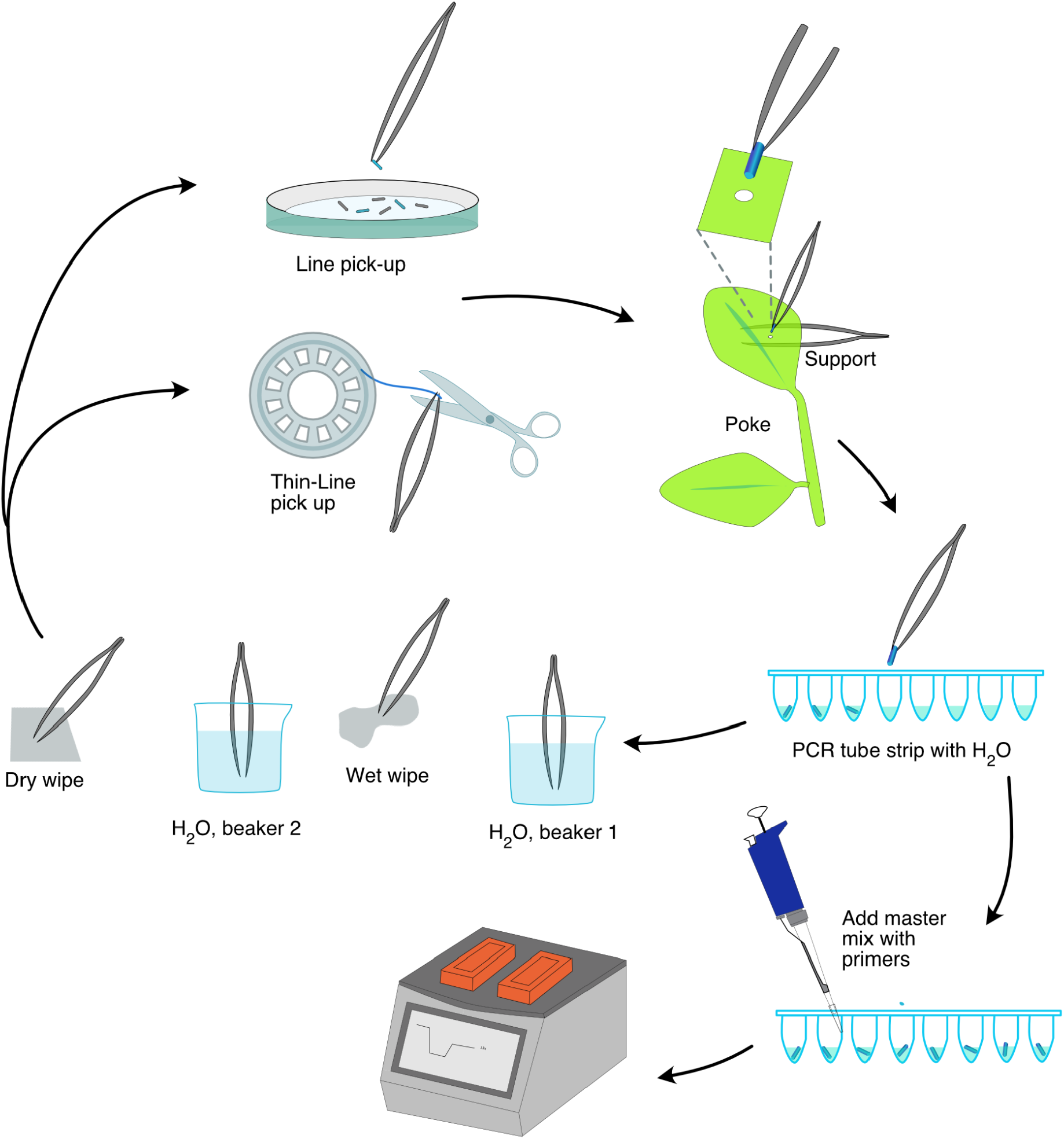
Line-PCR and Thin-Line-PCR workflow. Line (for example, 0.279 mm. diameter) can be pre-cut and picked up from a dish. Thin-Line (<0.1 mm. diameter) is easier to grasp and cut from the spool in the moment before the poke.

## Discussion

When choosing a PCR method, a researcher’s foremost concern might be accuracy. This would be the case, for example, in CRISPR screens in plants where false positives for the desired change and false negatives for the presence of the *cas9* gene can compromise a project. Another concern is cost and practicality of high throughput screens. To help address these concerns, we explored published and new variations of Direct PCR. We report practices that are compatible with high throughput studies and that at the same time maximize accuracy.

Line-PCR and CutTip have additional benefits. As compared to CTAB and similar DNA purification methods, Line, Thin-Line and CutTip improve sustainable chemistry. Isolation of DNA template with 0.279 mm. diameter line uses >99.9% less plastic than CTAB-like methods and 90% less plastic than NT-CutTip. These direct PCR methods also do not require hazardous chemicals that are commonly used in DNA purification, such as CTAB, chloroform and SDS. These Direct PCR methods require no modification to the PCR reagents. While some other probing materials that we tested such as glass, plastic, and metal jewelry beads had a sensitivity of zero, every line that we have tested yielded a high sensitivity. This is fortunate because an inexpensive spool of line is enough for thousands of reactions. Thin line is more expensive (∼$0.01 per reaction) and requires more labor than normal line, but thin line causes by far the least damage. Nevertheless, normal line (∼$0.001 per reaction) causes less damage than p20-based probes and traditional DNA purification methods. Importantly, Line-PCR is profoundly easy. The most difficult step is to wipe the tweezer with a paper towel between pokes, which is also important in traditional DNA purification. While we did not find it necessary to wear gloves while poking, we do recommend wearing simple eye protection. All of the poking was done at room temperature and seemed stable in room temperature water in the PCR tubes for several hours. This general easiness should enable PCR preparation in the field, with the caveat that Line in PCR reagent mix had diminished PCR sensitivity after ∼2 hours at room temperature.

Given CTAB, Direct PCR and intermediate methods, there are many valid methods to prepare a PCR template. We demonstrate high accuracy of some of the simplest plant Direct PCR methods to date, specifically the B.M. method and two methods that we developed: CutTip and Line. The accuracy, ease and speed of CutTip, B.M. and Line enable genotyping more plants and, if desired, more locations on a plant. Overall, quickly and confidently genotyping plants should improve the prospect of experiment completion.

## Data Availability

Details of each PCR are in Supplemental Tables 1 and 2.

## Acknowledgements

This work was funded by an Innovative Genomics Institute grant and an NSF-Plant Genome IOS Grant 144612: Rapid and Targeted Introgression of Traits via Genome Elimination. P.G.L. was supported by an NSF Graduate Research Fellowship, Elsie Taylor Stocking Fellowship and Eric and Louise Conn Graduate Student Award in Plant Biochemistry. We thank Nimrat Kaur for video editing. Maria Ximena Anleu Gil for contributing figure art and methodological insights, Thu Nguyen, Yinuo Zhang, and Varun Viswanath for technical support and Isabelle Henry for discussion.

## Author contributions

P.L. and L.C. wrote the manuscript, made figures, acquired funding, analyzed the data and supervised the research. P.L. and P.O.K. developed methodology and curated data. P.L., P.O.K., B.W. and E.M. performed the experiments.

