## Supplemental Figures for "Accurate Direct PCR with *Arabidopsis* and rice"

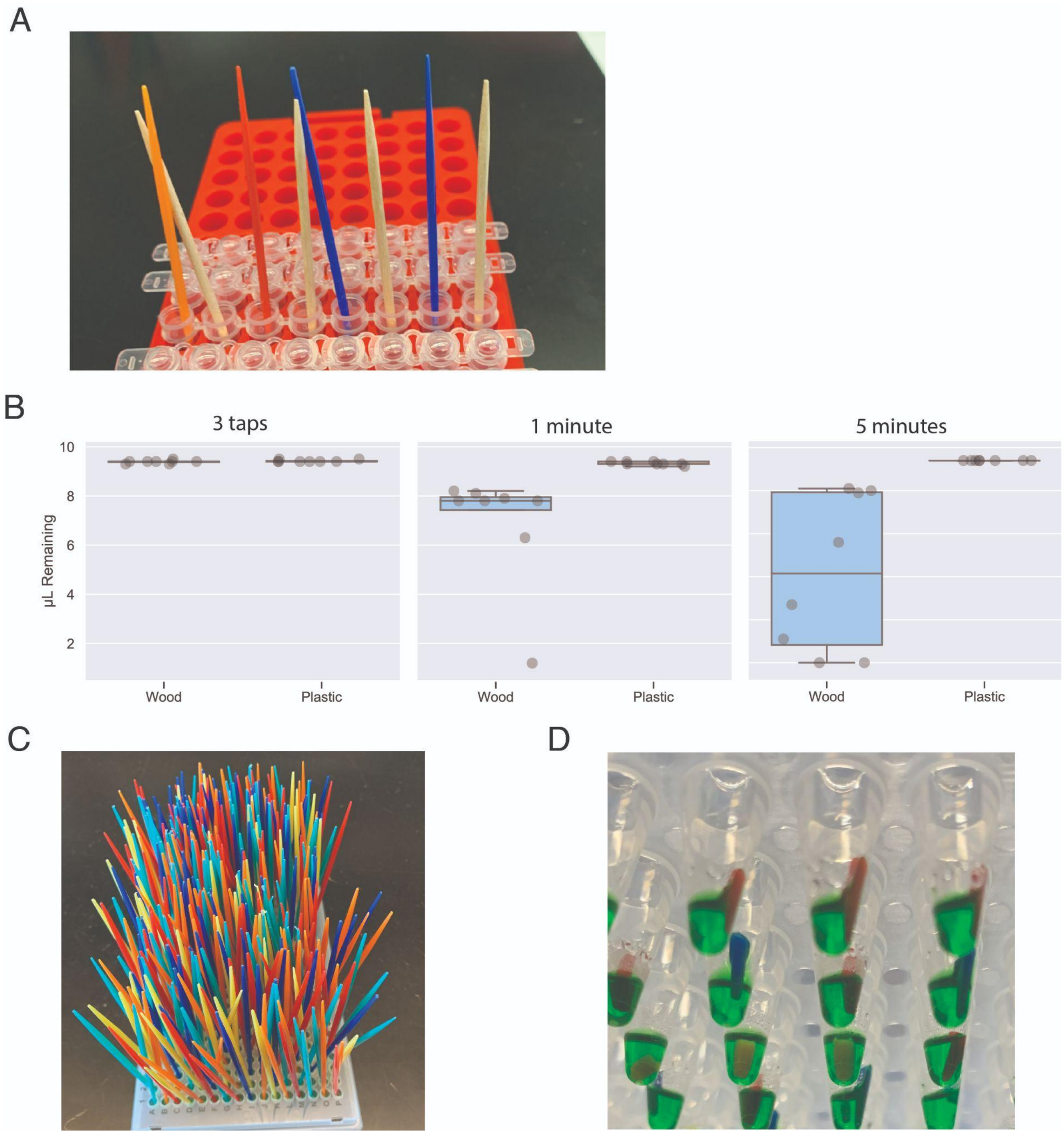


**Supplemental Figure 1**. Toothpicks for Direct PCR. A. Alternating wooden and plastic toothpicks placed into 10 µL of PCR reagents. B. Liquid remaining after: 1) Immediately tapping the tip on the bottom of the tube 3 times; 2) Sitting for 3 minutes then removing the toothpicks; 3) Sitting for 5 minutes then removing the toothpicks. C. Plate of 384 plastic toothpicks that were each poked into a leaf, then placed into 10 µL of PCR reagents. D. Testing cutting the tip of pipette tips, dropping them into 10 µL of PCR reagents where they stay. The plastic toothpicks were too difficult to cut.

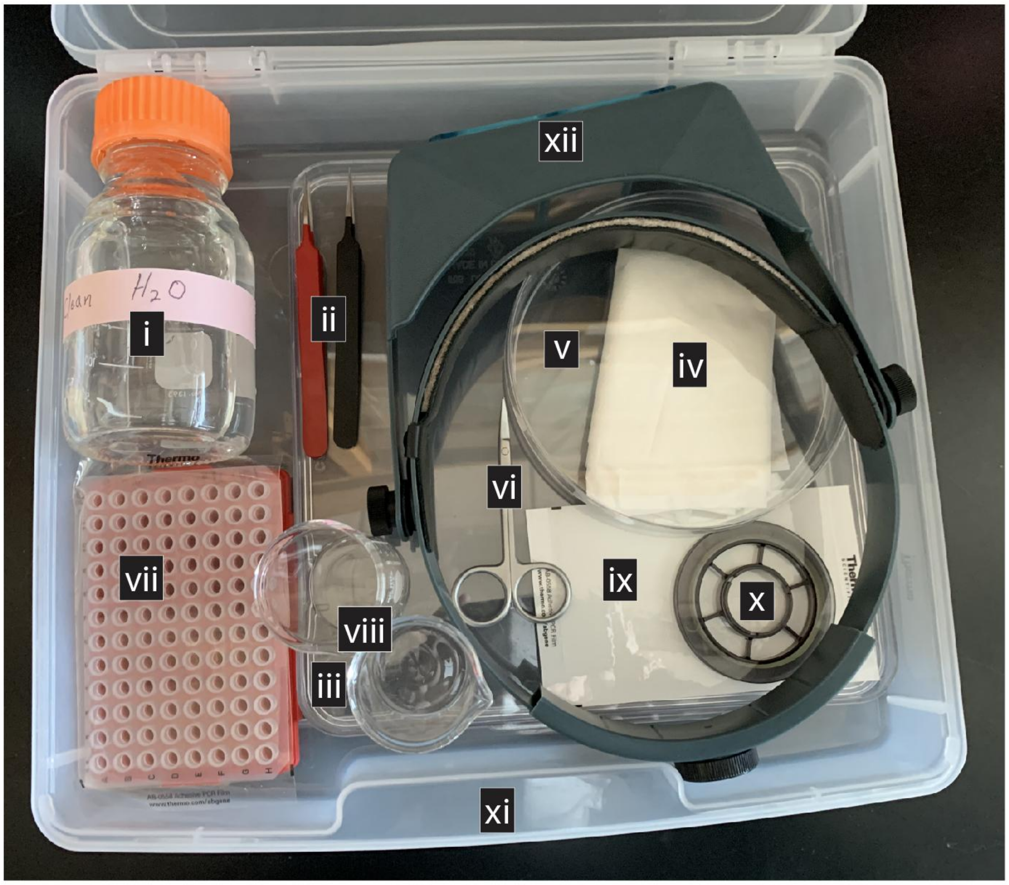


**Supplemental Figure 2**. Example kit for Thin Line-PCR. i. Clean water. ii. Two tweezers. iii. Large clean dish (to help prevent exposed thin line from touching contaminated surfaces) iv. Paper towels. v. Clean dish (for paper towels). vi. Scissors. vii. Sealed PCR tubes containing water or liquid reagents (not stored with kit). viii. Two beakers. ix. PCR sealing tape or caps. x. Spool of thin line (0.074 mm. diameter). xi. Container for kit. xii. Magnification / eye protection (such as OptiVISOR).

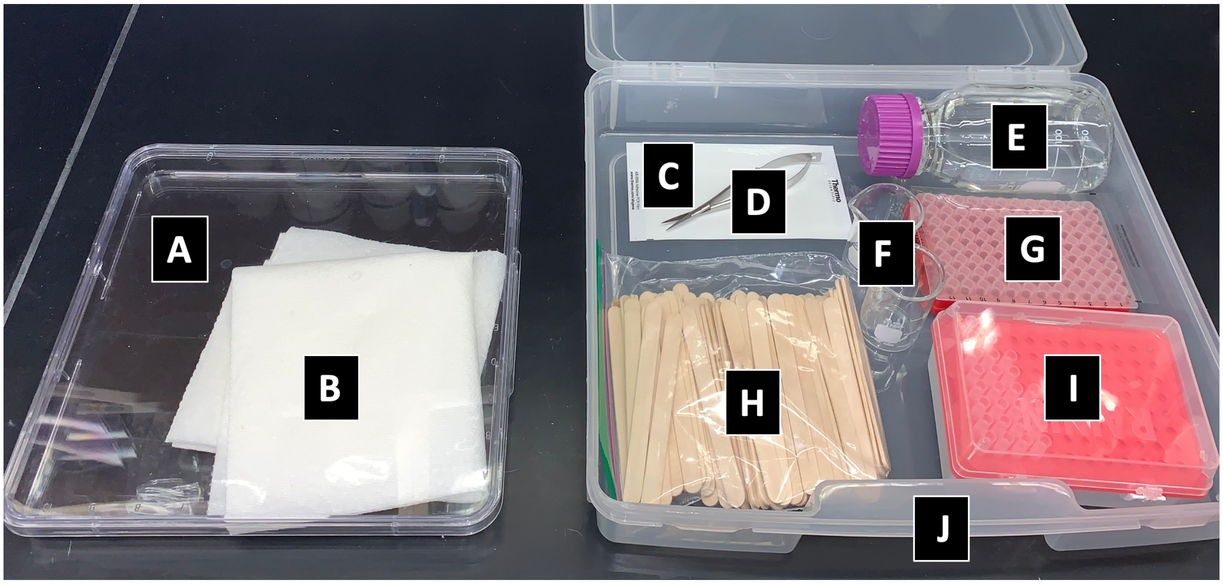


**Supplemental Figure 3**. Example kit for CutTip PCR. A. Clean dish. B. Paper towels. C. PCR sealing tape or caps. D. Scissors. E. Clean water. F. Two beakers. G. Sealed PCR tubes containing all liquid reagents (not stored with kit). H. Popsicle sticks (flat wooden toothpicks also work). I. p20 pipette tips. J. Container for kit.

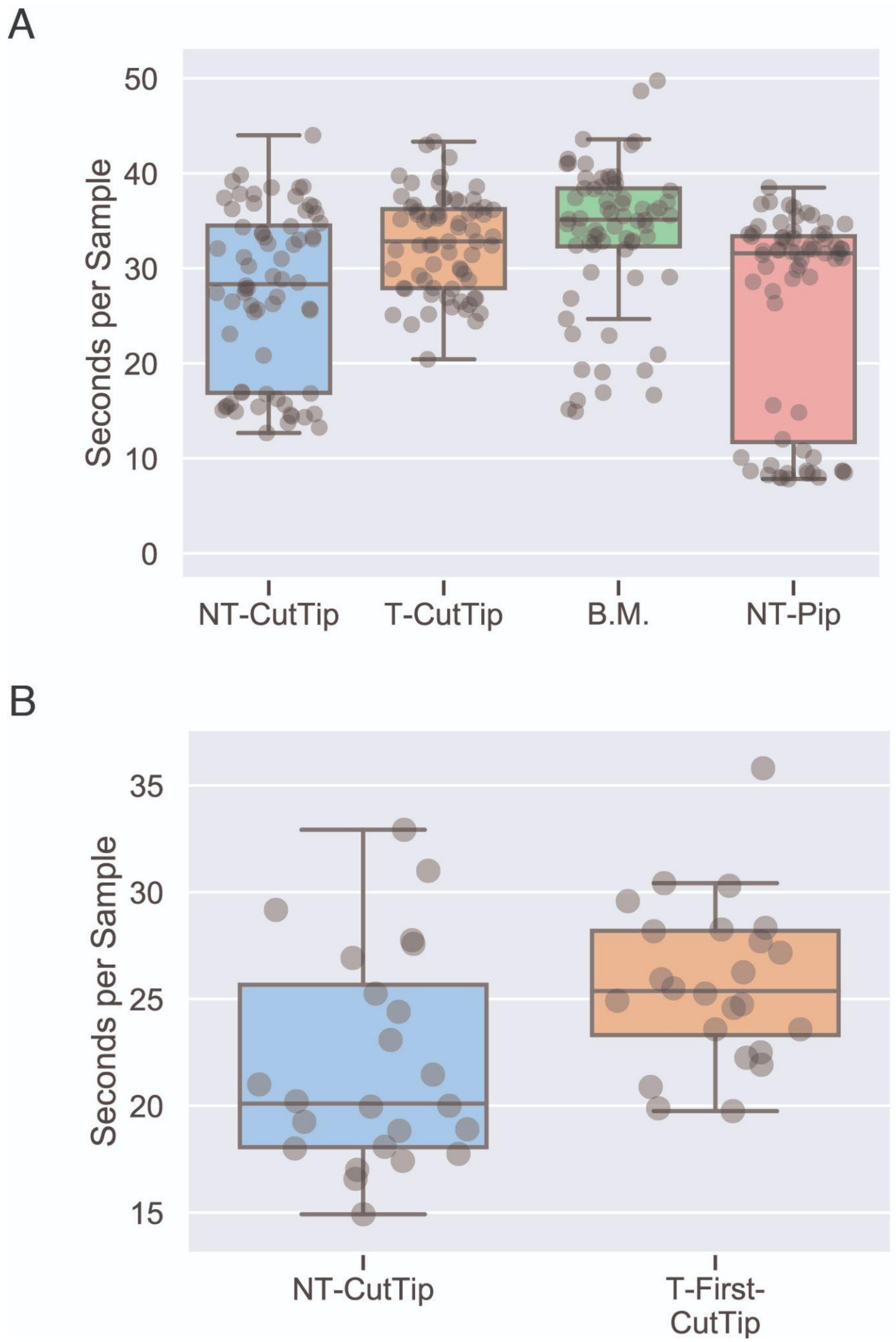


**Supplemental Figure 4**. Time to prepare a single sample, measured as the average number of seconds between 2 pokes in a series of 12 pokes. A. Comparison of 4 Direct PCR methods. Bimodal distributions can be explained by different technicians carrying out the procedure at different speeds, most obviously seen with NT-Pip. B. Comparison of 2 Direct PCR methods.

| Problem | Possible Reason | Suggested Solution |
|---|---|---|
| False Positives | DNA from previous samples reaching above cut. | Cut 2-3 mm. pieces. When poking, don't let tissue touch 1 mm. above bottom of tip. If tissue touches 1 mm. above bottom of tip, discard pipette tip. |
| | DNA from previous sample on scissors. | Use wet paper towel to wipe the entire scissor cutting surface. Keep paper towels on a non-contaminated surface. Use 2 beakers of cleaning water for dipping scissors as described. |
| | Other contaminated tools or reagents. | Use new reagents and cleaning water. Use filtered pipette tips when pipetting reagents into PCR plate (filter not necessary for poking for CutTip). Regularly clean all tools and surfaces. Use new popsicle stick or single-use flat wooden toothpicks. |
| | Non-specific primers. | Use 25-32 bp primers. Increase annealing temperature. |
| | CutTip hopped into wrong well. | Angle the tip into the well as seen in Figure 1.3C and Supplemental Movie. Don't cut above top of well. Iterative cuts are heavier, helping drop into correct well. |
| | Cut into wrong well. | Put informative stickers over plate near where cutting. Can validate by using a microscope to count one tip per well. |
| False Negatives | Tissue too large. | Use NT-CutTip, B.M. or T-First-CutTip. |
| | No template contacting PCR solution. | When poking, swivel the tip to get cell smear all around. If using visible tissue, make sure the tissue is not stuck in the pipette tip nor on the tube wall. |
| | Very faint band. | Increase PCR cycles. |
| | Several hours of sampling at room temperature. | If >2 hours of PCR preparation, cut tips into water. Add master mix within a few minutes of starting the thermal cycler. |
| | Mismatches between primer and target. | Design primers based on correct DNA sequences. Design primers based on Sanger sequencing the primer region using the plant of interest. |
| | Cut tip hopped into wrong well. | Angle the tip into the well as seen in Figure 1.3C and Supplemental Movie. Don't cut above top of well. Iterative cuts are heavier, helping drop into correct well. |
| | Cut into wrong well. | Put stickers on plate sealing tape near where cutting. Can validate using a microscope to count one tip per well. |

| | | |
|---|---|---|
| | Amplicon size too small or large. | Aim for 160-600 bp amplicons. |

**Supplemental Table 3**: CutTip Troubleshooting Guide

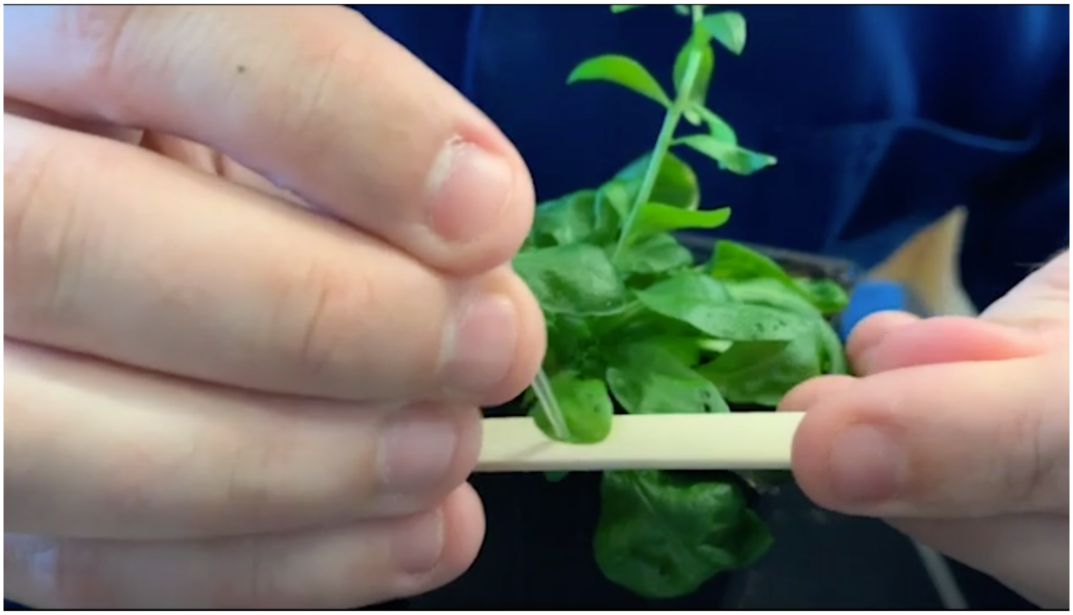

**Supplemental Movie 1**: Protocol of NT-CutTip PCR. Can be viewed at https://youtu.be/F-KiL9epodM

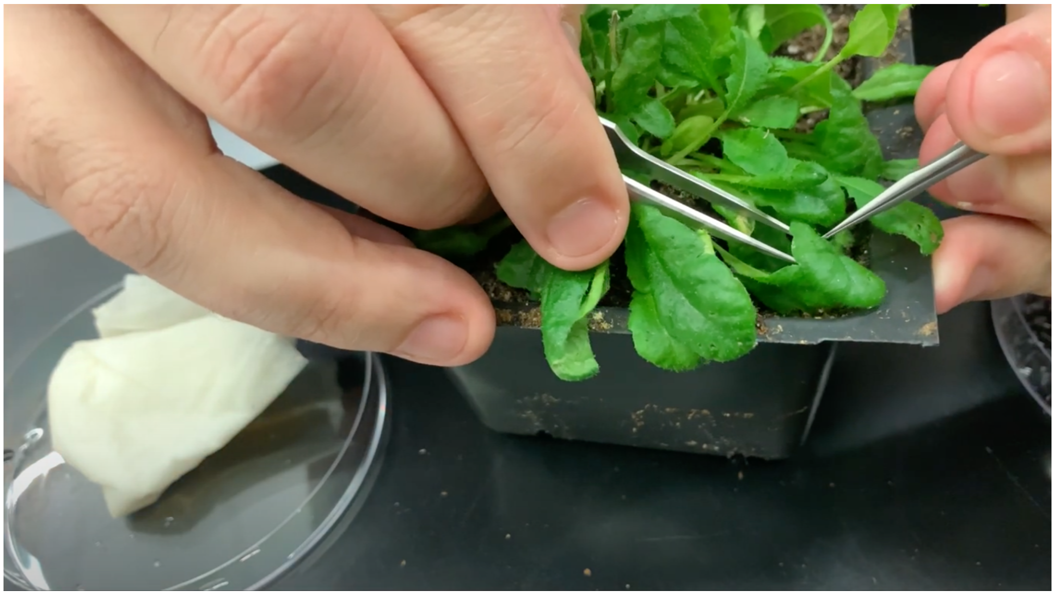

**Supplemental Movie 2** Protocol of Line-PCR. Can be viewed at https://youtu.be/Yy6DT8KVBQE
